# Volumetric Flow Imaging Microscopy to Enhance Particle Characterization

**DOI:** 10.64898/2026.06.23.733833

**Authors:** Farhan Khan, Benjamin Gincley, Farshid Khan, Ameet J. Pinto

**Affiliations:** School of Civil and Environmental Engineering, Georgia Institute of Technology, Atlanta, USA; School of Earth and Atmospheric Sciences, Georgia Institute of Technology, Atlanta, USA; Brook Byers Institute for Sustainable Systems, Georgia Institute of Technology, Atlanta, USA

## Abstract

Flow imaging microscopy (FIM) is an important technology for high-throughput characterization of microscopic particles and microorganisms. However, conventional FIM relies on single-plane imaging (SPI), resulting in out-of-focus particles, reduced measurement precision, and incomplete characterization of irregularly shaped objects extending along the z-axis. To address these limitations, a volumetric flow imaging (VFI) framework was developed and implemented on the portable ARTiMiS platform. This approach captures multiple frames along the z-axis and extracts the highest fidelity image for each particle, which can also be used for single image generation with all particles in focus (i.e., all in focus image) and for three-dimensional reconstruction of irregularly shaped objects. Benchmarking VFI with microspheres, live cells (*Chlorella vulgaris*), and filamentous cyanobacteria demonstrated increased fraction of particles in focus, reduced variability in particle size measurement, and increased resolvability of elongated particles in comparison to conventional SPI on commercially available FIM technologies. For *C. vulgaris*, VFI-derived size distributions closely matched curated FlowCam measurements without requiring post-processing to exclude out-of-focus particles. All-in-focus image reconstruction enabled simultaneous visualization of particles distributed across multiple depths and consistently resolved a greater proportion of filamentous structures as compared to SPI. For *Aphanizomenon* sp., *Dolichospermum* sp., and *Planktothrix agardhii*, the SPI approach captured only 84%, 61%, and 58%, respectively, of the total filament length resolved by AIF reconstruction. Beyond image-based characterization, VFI enabled estimation of dynamic particle properties such as sinking velocity and mass density. Application of this framework to *C. vulgaris* cultures revealed distinct mass-density trajectories under nitrogen-replete and nitrogen-deplete conditions, with cell mass density increasing over time under nitrogen-replete conditions and decreasing under nitrogen deprivation. Collectively, these results establish VFI as a next-generation framework for FIM that expands its analytical capabilities beyond conventional morphometric characterization and provides new opportunities for single-cell-enabled environmental monitoring and biomanufacturing.

## Introduction

Flow imaging microscopy (FIM) combines the high-throughput quantitative ability of a flow cytometer and collects images that can be analyzed to identify biotic and abiotic objects. As a result, flow imaging microscopy is gaining popularity for microbial community monitoring, especially for microalgae and phytoplankton monitoring. In addition to established commercial platforms such as Amnis ImageStream^x^ MK II (Cytek® Biosciences), FlowCam (Fluid Imaging Technologies, Inc.), Micro-Flow imaging (Bio-Techne), a number of custom FIM systems have recently been developed (Collins et al., 2019; Deglint et al., 2018; Gincley et al., 2024; Hardison et al., 2019; Li et al., 2024; Pollina et al., 2020). Several of these platforms have focused on reducing instrument cost and complexity, thereby broadening access to FIM and helping democratize the technology. When integrated with modern deep learning frameworks, FIM can support automated taxonomic classification, morphotype identification, and high-throughput ecological monitoring (Chan et al., 2023; Ciranni et al., 2024; Kraft et al., 2022; Luo et al., 2018; Madkour et al., 2023; Orenstein et al., 2020; Pastore et al., 2020; Xu et al., 2022).

Despite these significant advantages, FIM has several inherent limitations. In most FIM systems, the depth of the flow channel exceeds both the optical depth and the size of the particles being imaged. As a result, a substantial portion of the imaged particles are out-of-focus (Zölls et al., 2013). While reducing channel depth can mitigate this issue, it constrains the size range of particles that can be accommodated and increases the risk of channel clogging. The clogging issue is exacerbated when analyzing heterogeneous samples, including mixed cultures or environmental samples. Furthermore, flow imaging systems are susceptible to the presence of background or stagnant particles (Camoying & Yñiguez, 2016; Marvin et al., 2019; Romero-Martínez et al., 2017), which can artificially inflate particle counts. Both stagnant and out-of-focus particles result in erroneous particle counts and microbial community composition, where applicable, especially for low-concentration samples. In our previous study (Khan et al., 2025), two algorithmic methods were developed to detect and discard out-of-focus particles and stagnant particles. Discarding stagnant particles does not result in any loss of information, as only repeatedly observed particles are discarded. However, out-of-focus particles, if discarded, can lead to significant underestimation of particle counts (Gincley et al., 2025). Accurate estimation of particle counts (and microbial community composition, where applicable) requires that both in-focus and out-of-focus particles are considered. Moreover, out-of-focus particles present significant challenges for downstream analyses, particularly in taxonomic identification and semantic feature extraction, due to reduced image fidelity and loss of morphological detail (Kurinomaru et al., 2023; Lee & Kim, 2004; Ripple & Hu, 2015; Zölls et al., 2013). Thus, while preserving the out-of-focus may improve total biomass estimates, it does not help with cell characterization.

In this study, we present volumetric flow imaging (VFI) as a strategy to overcome key limitations of conventional FIM. The VFI approach was implemented on the ARTiMiS platform (Gincley et al., 2024), and its performance was evaluated against single plane imaging (SPI) acquired with ARTiMiS, as well as other flow imaging instruments. The results demonstrated that VFI not only mitigates limitations associated with out-of-focus particles but also addresses major challenges in imaging particles that extend along the depth of the flow channel, such as filamentous microorganisms (Graham et al., 2018). All-in-focus (AIF) image reconstructed from the optical slices acquired through VFI enables simultaneous visualization and detection of both small particles and large filaments, all in one single composite image. In addition to these advances, VFI enables estimation of dynamic particle properties, including sinking velocity and mass density, parameters that have only rarely been measured in flow through imaging systems (Bach et al., 2012; Miettinen et al., 2024). Collectively, these advances extend FIM’s functional capabilities by improving measurement reliability, image quality, and analytical precision.

## Materials and Methods

### Materials for imaging

#### Microspheres

Particle resolvability, AIF image generation, and cell density estimation were validated using commercially available microspheres of known size and density. For size estimation and resolvability experiments, polystyrene microspheres with nominal diameters of 2 µm, 4.5 µm, 6 µm, 10 µm, and 15 µm and a specific gravity of 1.05 g/cm³ were procured from Polysciences, Inc. For validation of density estimation, 4 µm polymethyl methacrylate (PMMA) microparticles with a density of 1.22 g/cm³ were obtained from Sigma-Aldrich, and 4.8-5.8 µm polystyrene microspheres with a density of 1.07 g/cm³ were obtained from Cospheric LLC. Microspheres from Polysciences Inc. and Sigma-Aldrich were supplied in a suspension, while microspheres from Cospheric LLC were supplied as solids. Solid microspheres were brought to suspension following the protocol described by Cospheric LLC.

#### Laboratory cultivation of spherical microalgae and filamentous cyanobacteria

*Chlorella vulgaris* cells were obtained from Algae Research Supply and used as a model for spherical microalgae. *Aphanizomenon sp*. (CAWBG01), *Planktothrix agardhii*, and *Dolichospermum flos-aquae*, all filamentous cyanobacteria with single-stranded morphology, were obtained from the culture collection at Bowling Green State University and used as model filamentous cyanobacteria. Cultures were grown in polycarbonate, unbaffled, sterile vent-cap flasks at nominal flask volumes of 125 mL (VWR International, 89095-258) in the laboratory. *C. vulgaris* was cultivated in Bold 1NV medium, whereas the filamentous cyanobacteria were maintained in Jaworski medium. Once every two weeks, a 1:10 dilution was carried out for maintenance by wasting the existing culture and adding fresh media to the culture flask. Cultures were continuously agitated using an orbital shaker in an environmental chamber set at 25 °C equipped with RGB LED lights. LED lights were set at a 12 h:12 h day-night cycle, maintaining a photosynthetic active radiation (PAR) value of ∼80 µmol-photons/m^2^s near culture flasks.

#### Cultivation of *Chlorella vulgaris* to induce lipid-accumulation mediated cell density changes

To induce changes in cell density during cultivation, lipid accumulation was inducted through nitrogen deprivation by using a modified Bold 1NV medium with no nitrogen. *C. vulgaris* cells were pre-grown in standard Bold 1NV medium. Two flasks (C1 and C2) were seeded from the parent cultures in 25 mL of Bold 1NV media at a cell concentration of 3.2 × 10^6^ cells per mL. Cells were collected from the parent culture, mixed together, gravity settled, and the supernatant was discarded to increase cell concentration. From the concentrated *C. vulgaris* inoculum, two flasks (NR1, NR2) were seeded on day 0 at a higher concentration (1 × 10^7^ cells per mL). Fresh Bold 1NV medium was added to the NR1 and NR2 flasks on day 5 to compensate for higher cell density. To induce nitrogen-depleted conditions, two additional flasks (ND1 and ND2) with 25 mL of custom Bold 1NV media were seeded with cells from the cell aggregate at 1 × 10^7^ cells per mL cell concentration. Samples were collected from each flask on days 4, 5, 6, 10, 13, 17, and 20 to carry out mass density measurement.

### Hardware development

#### Optical system design and calibration for volumetric flow imaging

The ARTiMiS platform, previously described by Gincley et al. (2024), was redesigned to integrate VFI capabilities. The base ARTiMiS was equipped with two M12 lenses in reverse configuration, affixed to the camera sensor installed on a stepper linear actuator for intermittent focal plane adjustment. The VFI-enabled ARTiMiS (subsequently referred to as ARTiMiS VFI) includes only the tube lens fixed directly to the camera sensor, while the objective lens is mounted on a voice-coil motor (VCM) to enable controlled axial translation. In this arrangement, the objective and tube lens function as an infinity-corrected microscope system, in which the objective produces a collimated beam that is subsequently focused onto the sensor by the stationary tube lens. The objective can be moved in the z-axis with high speed and precision using the VCM. This new optical system allows focus adjustment on a per-image basis, which improves the quality of images. The VCM also enables lens movement in the z-axis with minimal to no movement in the x-y planes. This enables seamless scanning through the depth of the flow channel.

The voice coil is controlled using the Arducam fork of the libcamera library. The relative position of the lens is in the range of 0 to 12. Lens position can be adjusted with a step size of 0.01. To translate the lens position to actual distance, VCM was calibrated using a step pyramid (Figure S1) as described previously (Yao et al., 2024) and fabricated using Nanoscribe with a 10x lens at the Institute for Matter and Systems (IMS), Georgia Tech. The calibration tool consists of one central pyramid and 4 smaller pyramids at the four corners. The central pyramid contains five discrete height steps (20 µm, 10 µm, 5 µm, 2 µm, and 1 µm) and was used for step size calibration. The four corner pyramids, each containing two steps, were primarily used to level the camera-lens assembly. Using the 5 µm and 10 µm steps of the central calibration pyramid, the axial displacement of the VCM was calibrated, and a reported 0.01 increment in lens position corresponded to an axial movement of 1 µm. The camera-lens assembly was subsequently leveled using the 10 µm step of all 5 pyramids. ARTiMiS VFI can capture images at 1 µm intervals, but this generates a higher number of images, which increases the time required to do the volume scanning. Considering the objective lens having an approximate depth of field of ∼10 µm, a step size of 10 µm was selected as optimal to scan the volumetric field of view (VFOV) while minimizing scanning time.

### Software development

#### Particle detection in a volumetric scan

An image processing pipeline was developed to detect and extract particles from the VFI image stack. First, each image in the VFI-acquired image stack was denoised, and an initial binary mask was created using a contrast threshold. The binary masks were then processed through multiple morphological transformations to create the final binary mask. Three-dimensional connected component labeling was performed on the binary mask stack using the “label” function from the ‘scipy.ndimage’ module (SciPy v1.15.1) to label individual particles. For each labeled particle, sharpness was computed for each image slice using a Sobel filter implemented in scikit-image (v0.24.0), and the slice with the maximum sharpness was extracted for further downstream analysis.

#### All-in-Focus image generation and denoising

A computer vision-based algorithm was developed to generate AIF images from an image stack. For each image, a gradient map was computed using a Sobel filter. For each x-y pixel location, the pixel with the maximum gradient value along the z-axis was selected to construct an initial compound image. This compound image exhibited pixelation artifacts and therefore required post-processing. A lightweight U-Net–based convolutional neural network model was trained to denoise the AIF image generated from the z-stack. The model has an encoder and a decoder framework. Encoder and decoder are symmetrical, each with three UNetBlocks and three max pooling layers. A bottleneck block operating at the lowest spatial resolution captures high-level contextual information before reconstruction. The training dataset was created from the VFI dataset of microspheres, *C. vulgaris* cells, and filamentous cyanobacteria to incorporate a broad range of particle morphologies, sizes, and imaging characteristics. Ground truth patches were extracted from the most-in-focus (MIF) images in each volumetric scan. Patch size was selected to be 154 x 154 pixels for model training. Corresponding reconstructed AIF-derived image patches containing reconstruction-associated noise artifacts were obtained by identifying the spatially best-aligned crop within a 40-pixel search radius centered on the ground-truth patch location. Alignment was determined by minimizing the mean squared error (MSE) between candidate patches and the target ground-truth patch. The resulting paired images were then divided into training (2156 image pairs) and validation (380 image pairs) datasets. To improve model robustness and reduce overfitting, the training dataset was augmented using combinations of image rotation and flipping operations, including 45°, 90°, and 135° rotations, as well as horizontal and vertical flips. Through augmentation, each original image pair generated a total of 12 augmented image pairs.

#### 3D reconstruction from volumetric flow imaging

A computational pipeline was developed to generate three-dimensional representations of imaged structures within the volumetric field of view (VFOV) from VFI data using filamentous organisms as a case study. A custom masking method was developed based on the recommendation by Iamsiri et al. (2019). Each image in the volumetric image stack was converted to grayscale, the Tenenbaum gradient was measured at each pixel, and the gradient map was then normalized and denoised. A binary mask was generated from the denoised gradient map by thresholding. Multiple morphological operations (binary opening, remove small objects, binary closing, dilation, and erosion) were carried out on the binary mask. A ‘filled mask’ was generated from the binary mask by filling the enclosed area. The processed binary mask was then subtracted from the filled mask, and small holes were removed to get the final mask. Then, using the ‘label’ function from the ‘scipy.ndimage’ module, volumetric point clouds for each filament were generated from the final mask stack. Center axes of the filaments were determined by applying the ‘skeletonize’ function to the 3D point cloud. In tangled filament networks, intersections between multiple filaments can produce crisscrossing centerlines at junction regions. To isolate individual filament fragments, the skeletonized output was further processed using the ‘Skeleton’ function from the ‘Skan’ library (v0.13.1). Subsequently, a bipartite matching framework based on the linear sum assignment algorithm from ‘scipy optimize’ was used to reconstruct individual filaments from the fragments, prioritizing spatial proximity in x, y, and z dimensions, as well as angular alignment between segment endpoints. Filament width was measured along the center axes. Finally, filaments were represented as three-dimensional cylindrical structures, where the local filament width defined the cylinder diameter and the reconstructed centerline defined the cylinder axis.

### Evaluation framework and benchmarking methodology

#### Benchmarking ARTiMiS VFI against commercially available alternatives

To compare the size measurement accuracy of the newly developed method, ARTiMiS VFI was compared to commercially available flow imaging instruments, i.e., FlowCam VS (Fluid Imaging Technologies) and Amnis ImageStream^x^ MK II. For this evaluation, 4 µm, 10 µm, and 15 µm beads were imaged on all three devices. FOV80 flow cell (FC) with 10x objective was used on FlowCam to acquire image data. Visual Spreadsheet was used to process particle images and extract features. Image data was acquired using INSPIRE acquisition software and the 40x objective lens, and analyzed using IDEAS 5.0 software. Particle size (area) reported by IDEAS 5.0 was multiplied by a factor of 0.25 to get area in µm^2^. The diameter of the particle was calculated from the measured area using this equation:

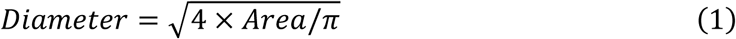

#### Image sharpness evaluation metric

Edge noise gradient, a gradient-based metric, was computed to quantify the sharpness of particle images. First, images were smoothed using a Gaussian filter (OpenCV v4.11) with a kernel size of 3 to reduce high-frequency noise while preserving structural edges. The smoothed image was then processed using a Sobel operator (OpenCV v4.11) to compute the first-order spatial derivative. Finally, the mean and standard deviation of the gradient image were calculated using the ‘meanStdDev’ function from the OpenCV library (v4.11). The standard deviation of quantified variability in edge intensity is reported as the edge noise gradient.

#### Measurement distribution comparison using Cohen’s d

To compare two distributions of measurements, Cohen’s d effect size was calculated. Cohen’s d quantifies the difference between the means of two populations relative to their pooled standard deviation. The magnitude of Cohen’s d reflects the degree of separation between the two distributions, while the sign indicates the direction of the difference. Cohen’s d was measured using the following equations:

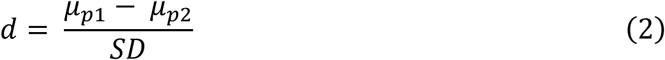

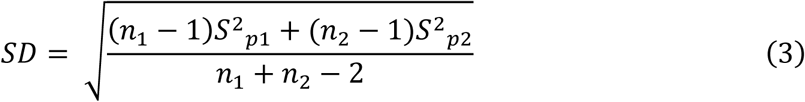

Where d = Cohen’s d, SD = pooled standard deviation, *μ_p_*_1_ = mean of population 1, *μ_p_*_2_ = mean of population 2, *n*_1_, *n*_2_ = population sizes, and *S_p_*_1_, *S_p_*_2_ = standard deviation of the two populations.

#### Assessment of filament resolvability in AIF images

Two metrics, total filament area and total filament length, were primarily used to compare filament detection in AIF images to that of most-in-focus (MIF) images. The MIF image was the image with the highest sharpness score within the volumetric image stack. AIF and MIF images were processed to get foreground masks following the same methodology described in the ‘3D reconstruction from volumetric flow imaging’ section. For the binary mask generation step, thresholding was performed at the 30th percentile of the gradient value for *Aphanizomenon* and *Dolichospermum* and the 10th percentile for *Planktothrix* and mixed filaments. *Planktothrix* filaments were subvisible, requiring a smaller threshold value to detect them (Figure S2). The final mask was skeletonized using the “skeletonize” function. The resulting skeleton had one major line representing the filament with small transverse lines. Transverse lines were pruned using the “prune” function from the plantcv library. Finally, the total length of the pruned skeleton was computed to measure the total length of all resolvable filaments.

#### Noise measurement for AIF images

Noise levels of images were measured using Median Absolute Deviation (MAD) of the high-frequency residual image (Khalil et al., 2008). Images were converted to greyscale and smoothed using the uniform filter from the ‘scipy.ndimage’ library. Image noise was estimated by subtracting the smoothed image from the original grayscale image to isolate high-frequency residual components. The residual noise level was then quantified using the MAD, scaled by a factor of 0.6745, and reported as the noise level.

#### Estimation of dynamic particle properties

Two volumetric scans of the same field of view (FOV), separated by a short and controlled time interval, can provide a direct measurement of particle displacement along the z-axis with respect to time (Figure S3). From the vertical settling distance and the elapsed time between scans, settling velocity was computed for each particle. For each FOV, two volumetric scans were carried out from the bottom to the upward direction. The z-axis position and timestamp of each optical section were recorded. Within each volumetric scan, the z-position of individual particles was estimated from the optical section in which the particle exhibited maximum sharpness. Particles detected in the two scans were subsequently paired using a bipartite matching algorithm to establish particle correspondence across scans (Figure S3). The two z-axis positions associated with each particle from consecutive volumetric scans were used to determine the settling distance (*s*), while the difference in acquisition timestamps provided the settling duration (*t*). Particle velocity (*v*) was measured using the following formula:

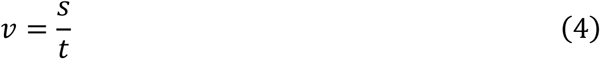

Theoretical sinking velocity (*v_t_*) of a particle was computed using Stokes’ law:

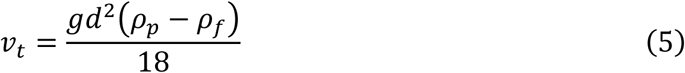

where acceleration due to gravity, g = 9.79506 m/s^2^, dynamic viscosity of fluid, *ρ_f_*= 1000 kg/m^3^. Density of particle, *ρ_p_* and the diameter of the particle, d, was acquired from the supplier’s catalog.

The measured sinking velocity was used to estimate particle mass density using Stokes’ law as follows:

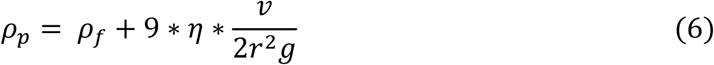

Where g is gravitational acceleration, *η* is dynamic viscosity (1.0016×10^-3^ kg/ (m.s)), *ρ_f_* is the mass density of fluid (1000 kg/m^3^) and *ρ_p_* is the mass density of particles in kg/m^3^.

## Results and Discussion

### VFI enhances particle resolvability, focus quality, and characterization accuracy

A single plane image (SPI), whether acquired with ARTiMiS or FlowCam VS, cannot detect all the particles within the VFOV. While allowing settling time prior to imaging with ARTiMiS can be used as a mitigation strategy, the settling time required for different particles can vary. As a result, SPI preferentially captures particles that settle faster and into the chosen focal plane, ultimately skewing sample characterization. To overcome this, we employed the VFI method. VFI was compared against SPI using microspheres with diameters of 2 µm, 4.5 µm, 10 µm, and 15 µm and a density of 1.05 g/cm^3^. Single plane imaging was performed with a short settling time (15 seconds), and volumetric scanning was performed without settling. Detected microspheres were sorted as in-focus particles, out-of-focus particles, and aggregates. Aggregates are instances where multiple microspheres appear in the same region of interest (ROI) and were excluded from downstream analysis as automated feature measurement on particle aggregates is not reliable. Figure 1A presents percentages of in-focus, out-of-focus, and aggregates in SPI and VFI derived microsphere population with different sizes. Counts for each category were normalized by the highest total count for each microsphere size. For smaller microspheres (2 µm and 4.5 µm), VFI captured a greater portion of microspheres in-focus than SPI. SPI was only able to capture 61.3% and 52.5% of the microspheres in-focus for the 2 µm and 4.5 µm populations, respectively. Approximately 16.2% of the detected 4.5 µm microspheres were out-of-focus, and this fraction increased to 23.2% for 2 µm microspheres.

**Figure 1:**
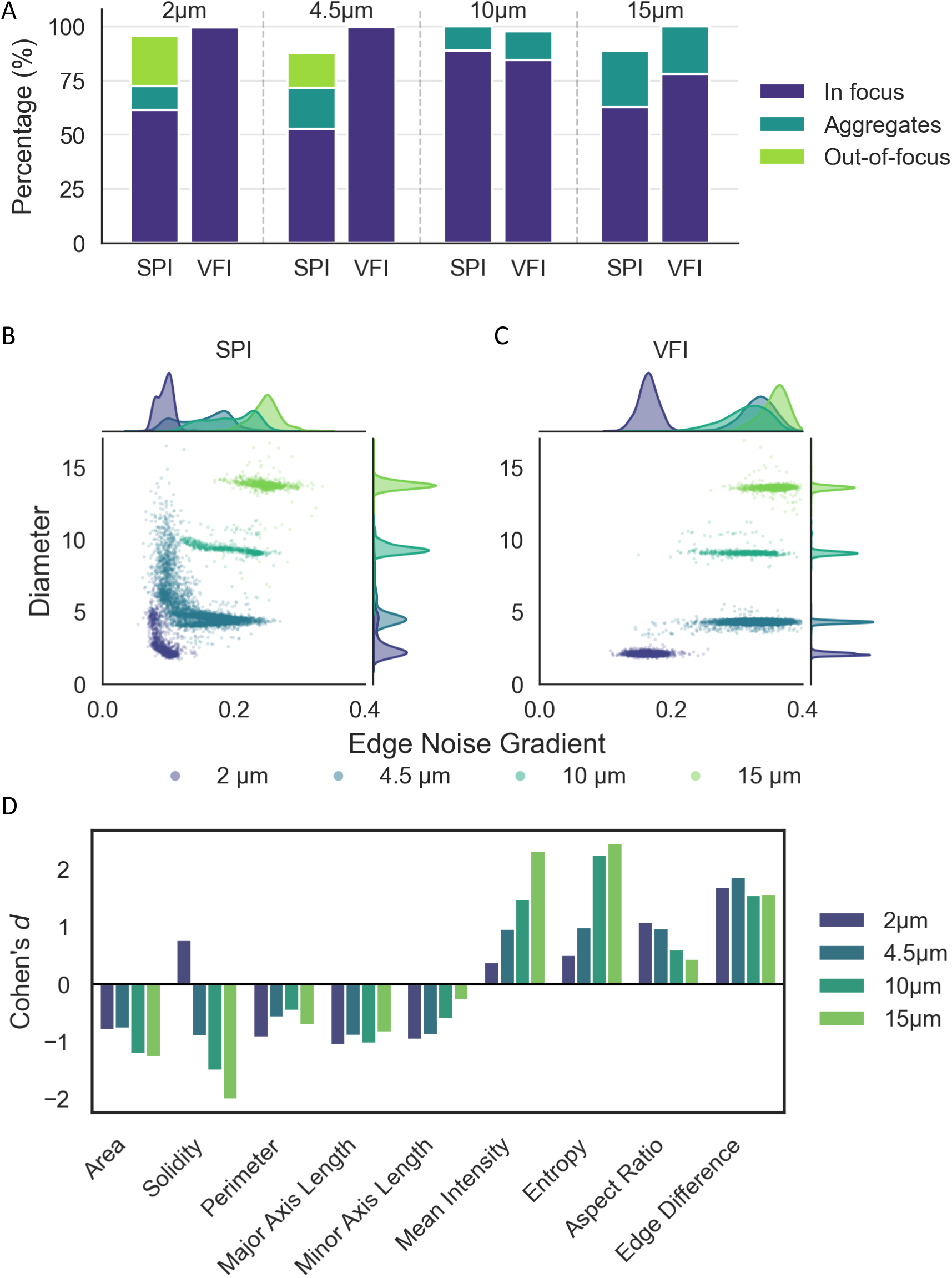
(A) Percentage of in-focus, aggregated, and out-of-focus particles detected using single-plane imaging (SPI) and volumetric flow imaging (VFI) for 2, 4.5, 10, and 15 µm microspheres. Percentages are normalized to the higher total particle count obtained by either method for each particle size. (B–C) Scatter plots of particle diameter versus edge noise gradient for microspheres measured using SPI (B) and VFI (C). (D) Cohen’s d comparing feature distributions between SPI and VFI; positive values indicate higher feature values for the VFI method.

Representative microsphere ROIs acquired using SPI and VFI are provided in the Supplementary Information (Figure S4). This behavior is expected, as smaller particles settle more slowly and therefore remain suspended across a broader range of depths within the imaging volume. For larger spheres (10 µm and 15 µm), both the SPI and VFI methods had no particles out-of-focus as these particles settle faster than smaller particles. SPI detected a marginally higher proportion of in-focus 10 µm microspheres (approximately 5% greater than VFI), whereas VFI detected 19.5% more in-focus 15 µm microspheres relative to SPI. This discrepancy may be associated with differences in the extent of particle aggregation between the samples.

The binary classification of particles as either in-focus or out-of-focus does not fully capture the optical reality of image formation, as focus is inherently a continuous rather than a discrete phenomenon. When a particle is visible but out of the depth of field, the particle appears blurry, and structural details are diminished. The degree of detail loss increases with axial distance from the optimal focal plane. To quantify the blurriness, we used the edge noise gradient parameter. Figure 1B and 1C show the edge noise gradient of particles detected by the SPI and VFI method, with a high edge noise gradient indicating a sharp transition, while a lower gradient value is associated with blurry images. For each size of microspheres, VFI-derived particle images scored a higher edge noise gradient than SPI (Table 1). Statistical analysis indicated that the differences were highly significant for all particle sizes (p < 0.001). Thus, multiplane acquisition of the VFI method mitigates detail loss due to defocus by incorporating information across multiple focal depths, thereby improving overall image sharpness and particle resolvability.

**Table 1:** Comparison of edge noise gradient between SPI and VFI methods across different particle sizes.

| Microsphere Size | Mean edge noise gradient (SPI) | Mean edge noise gradient (VFI) | Increase in VFI(%) | *p-value* |
| --- | --- | --- | --- | --- |
| 2 $\mu\text{m}$ | 0.01 | 0.17 | 73.68 | <0.001 |
| 4.5 $\mu\text{m}$ | 0.15 | 0.33 | 114.28 | <0.001 |
| 10 $\mu\text{m}$ | 0.19 | 0.31 | 61.85 | <0.001 |
| 15 $\mu\text{m}$ | 0.28 | 0.36 | 44.53 | <0.001 |

SPI not only results in lower edge noise gradient, but also increases deviation from the true diameter, especially for 2 µm and 4.5 µm particles (Figure 1B and 1C), relative to the VFI approach. Particles derived using VFI maintain a higher edge noise gradient and a narrower distribution of measured diameters than respective SPI-derived particles (Table 2). Variance and coefficient of variance (CV) were lower for VFI than for SPI across all microsphere sizes. Bias represents the difference between the average measurement and the standard size specified by the manufacturer. Both VFI and SPI had comparable negative bias (<10%) for 10 µm and 15 µm microspheres. This negative bias likely resulted from the light refraction effect and the artifact from the LED assembly (Figure S4). For smaller particles (2 µm and 4.5 µm), the refraction effect and the LED assembly artifact were minimal, and the edges of the microspheres were continuous. VFI exhibited minimal measurement bias for these microsphere populations, with bias values of 0.05 and -0.04 for the 2 µm and 4.5 µm microspheres, respectively. These results highlight the remarkable accuracy and robustness of the VFI method over the SPI. To compare VFI and SPI methods with respect to additional semantic features measured by ARTiMiS, Cohen’s d effect size was calculated for each feature distribution. Lower absolute values of Cohen’s d indicate that the distributions of a feature acquired by VFI and SPI are similar, and a higher value signifies divergence. The sign of Cohen’s d denotes the direction of the difference: negative values indicate higher feature values measured using SPI, while positive values indicate higher feature values measured using VFI. We observed a positive Cohen’s d value for mean intensity, entropy, aspect ratio, and edge difference. Higher mean intensity corresponds to a brighter image, while higher entropy indicates higher information. Increased edge difference signifies sharper edge transition (sharper image), and a higher aspect ratio (minor axis length/major axis length) means higher circularity. These features indicate a better quality of ROI extraction and possibly better feature extraction. Negative Cohen’s d values were found for area, solidity, perimeter, major axis length, and minor axis length. This indicates that features derived from SPI were greater than those of the VFI method. As particles often appear larger due to out-of-focus, negative Cohen’s d for these features were expected.

**Table 2:** Statistical analysis of microsphere size measurement using the VFI and the SPI methods.

| Microsphere Size | Var (VFI) | Var (SPI) | CV% (VFI) | CV% (SPI) | Normalized Bias (VFI) | Normalized Bias (SPI) |
| --- | --- | --- | --- | --- | --- | --- |
| 2 $\mu\text{m}$ | 0.01 | 1.35 | 5.42 | 40.45 | 0.05 | 0.44 |
| 4.5 $\mu\text{m}$ | 0.02 | 2.43 | 2.82 | 30.06 | -0.04 | 0.15 |
| 10 $\mu\text{m}$ | 0.08 | 0.26 | 3.18 | 5.47 | -0.09 | -0.06 |
| 15 $\mu\text{m}$ | 0.07 | 0.47 | 1.87 | 4.97 | -0.09 | -0.08 |

For some of the features, we also observed a gradual change in Cohen’s d with a change in the size of the microspheres. Larger microspheres exhibit higher settling velocities and therefore reach the focal plane more rapidly than smaller particles. As a result, differences between VFI and SPI became progressively reduced for larger microspheres, and the absolute value of Cohen’s d for minor axis length and aspect ratio decreased as the size of the microspheres increased. However, mean intensity and entropy had opposite trends. VFI-derived ROIs had higher light intensity and had more information with an increase in microsphere size in comparison to SPI-derived ROIs. Thus, VFI may have a smaller advantage over SPI in terms of size estimation for larger particles, but its advantage in terms of resolving structural information is high for larger particles.

### ARTiMiS VFI demonstrated more precise particle characterization as compared to commercial flow imaging instruments

Errors in particle size measurements arising from out-of-focus imaging have been reported for commercial flow imaging systems. Multiple studies have reported that optical defocus can introduce substantial inaccuracies in semantic feature extraction and morphometric measurements (Kurinomaru et al., 2023; Ripple & Hu, 2015; Zölls et al., 2013). To evaluate the impact of defocus on particle size estimation and to benchmark the performance of ARTiMiS VFI, standardized microspheres with nominal diameters of 4.5 µm, 6 µm, and 10 µm were imaged and analyzed on ImageStream^x^ Mark II, FlowCam VS, and ARTiMiS VFI. Figure 2A presents the distribution of measured diameter for each microsphere population across the three imaging platforms. The FlowCam used for this study required manual focusing before the sample run. Once a sample run is in progress, focus cannot be adjusted as adjusting focus during a sample run disrupts FlowCam’s background compensation, producing background artifacts that compromise statistical accuracy. Fixed focus mode of FlowCam resulted in two distinct peaks in the measured size distributions for all three microsphere populations (Figure 2A), which agrees with the findings of Zölls et al. (2013). The first peak (4.6 µm) corresponds to particles imaged in focus, whereas the second peak, characterized by larger measured diameters (5.3 µm), represents out-of-focus particle populations. For 4.5 µm microspheres, the first peak was at 4.6 µm and the second peak at 5.3 µm with a median of 5.58 µm. The bimodal distribution is nearly symmetric, with the out-of-focus mode accounting for almost half of the total data (49%). The percentage of out-of-focus mode increased for 6 µm microspheres (58.5%) and decreased for 10 µm microspheres (46%). Because of the bimodal distribution, interquartile ranges (IQR) were the highest among the three instruments for each microsphere size (0.689, 1.341, and 1.437 for 4.5 µm, 6 µm, and 10 µm microspheres, respectively).

**Figure 2:**
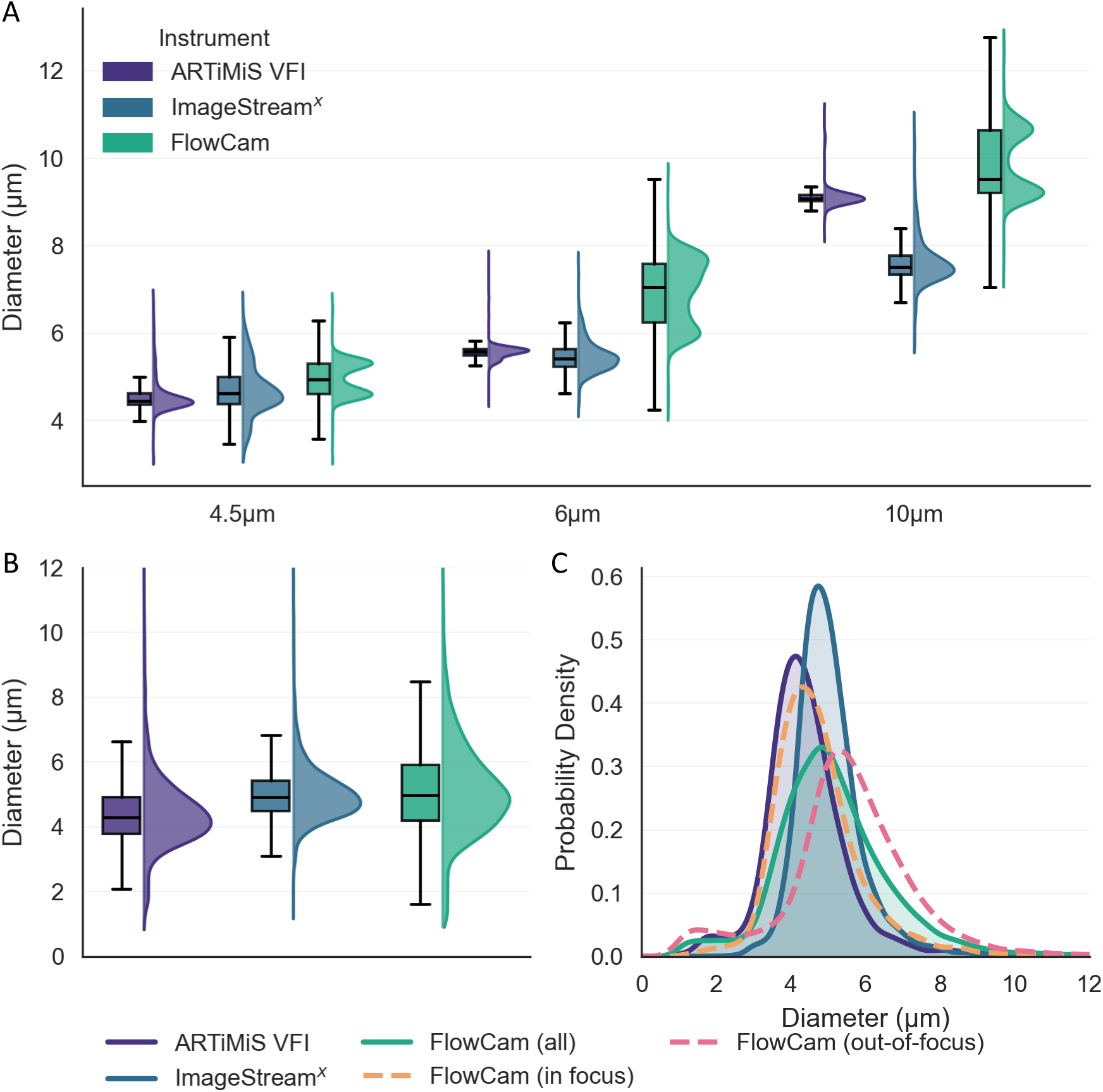
(A) Half-violin plots of particle diameter for standard microspheres measured using ARTiMiS VFI, Amnis ImageStream^X^ MK II, and FlowCam. The ARTiMiS-derived distribution is narrower compared to the other instruments, indicating improved precision. The FlowCAM distribution exhibits two distinct peaks corresponding to in-focus and out-of-focus particle populations. (B) Half-violin plots of C. vulgaris cell diameters measured using the same instruments. ImageStream^X^ shows the narrowest distribution, while FlowCAM exhibits the widest. Unlike the microsphere measurements, the FlowCAM distribution for C. vulgaris does not display clearly separated peaks. (C) Density distributions are further shown for ARTiMiS, ImageStream^X^, and FlowCam (combined, in focus, and out-of-focus subsets). For FlowCam, the in-focus and out-of-focus populations exhibit distinct peaks, consistent with the microsphere results, with the in-focus distribution closely matching that obtained from ARTiMiS.

ImageStream^x^ has active autofocus capabilities. It synchronizes the camera to the sample flow rate and continuously focuses on the particles using SpeedBeads, which serve as an internal control. Because of the active autofocus, ImageStream^x^-derived size measurements had single peaks and long tails (IQR = 0.613, 0.407, and 0.432 for 4.5 µm, 6 µm, and 10 µm microspheres, respectively). In comparison, the ARTiMiS-derived measurement showed single narrow peaks with the smallest IQRs among the three instruments (IQR = 0.252, 0.155, and 0.144 for 4.5 µm, 6 µm, and 10 µm microspheres, respectively), indicating high precision. This demonstrates that VFI not only improves particle resolving capacity, but it also provides better particle characterization than commercially available expensive instruments.

To evaluate the performance of the three imaging platforms under realistic biological conditions, *C. vulgaris* cells obtained from the same culture flask were analyzed using all three instruments. Unlike standardized microspheres, cell cultures exhibit inherent size heterogeneity, providing a more representative test of particle characterization performance. Consequently, this experiment enabled direct comparison between volumetric flow imaging (ARTiMiS VFI), autofocus-enabled single particle imaging (ImageStream^x^), and fixed focus single plane imaging (FlowCam). ARTiMiS VFI yielded a lower mean measured diameter (4.45 µm) compared to ImageStream^x^ (5.06 µm, p < 0.001) and FlowCam (5.16 µm, p < 0.001), with corresponding standard deviations of 1.28, 1.03, and 1.67 µm and IQR of 1.14, 1.72, and 0.94, respectively. ARTiMiS VFI-derived size distribution diverged from both FlowCam (d = 0.46) and ImageStream^x^ (d = 0.53). ImageStream^x^ utilizes SpeedBeads for autofocus calibration, and the thresholding procedure used to exclude these beads from analysis also removed a portion of smaller particles within the sample matrix. This likely contributed to the underrepresentation of particles below 3 µm and the corresponding upward shift in the measured size distribution and narrower IQR. For FlowCam, separation of the particle population into in-focus and out-of-focus subsets using an edge gradient (from Visual Spreadsheet) threshold (edge gradient > 50 for in-focus and < 50 for out-of-focus particles) revealed two distinct density peaks. The in-focus population exhibited a mean diameter of 4.78 µm and an IQR of 1.28, whereas the out-of-focus population exhibited a larger mean diameter of 5.57 µm and an IQR of 1.7. The in-focus FlowCam distribution closely matched the ARTiMiS-derived distribution (mean = 4.45 µm, IQR = 1.14), resulting in a substantially lower Cohen’s d value of 0.244. Collectively, these findings suggest that ARTiMiS VFI can accurately estimate biological cell size distributions without requiring additional post-processing or filtering to exclude out-of-focus particle populations.

### Volumetric Flow Imaging Enables Complete Filament Resolution and Three-Dimensional Orientation Reconstruction

As demonstrated in the preceding section, spherical particles ranging from 2 to 15 µm can be captured near their optimal focal plane using VFI, enabling the extraction of each particle from its corresponding focal depth. However, each optical section contained a subset of the total particle population within the VFOV. AIF image reconstructed from the volumetric scan integrates particles across all focal planes into a single composite image, providing a more comprehensive representation of the particle population (Figure S5). The utility of AIF imaging becomes particularly pronounced for the analysis of particles such as filaments that extend across multiple axial planes along the z-axis. Even the VFI method cannot resolve filamentous objects fully, as often no single plane contains full information about the filaments. Fully resolving filamentous objects, therefore, requires combining image data across multiple focal planes, a capability absent in existing flow-imaging systems, making filamentous objects particularly challenging to characterize (Graham et al., 2018). To address this, volumetric scans of *Aphanizomenon* filaments were acquired using ARTiMiS VFI with a 10 µm step size, where each optical section captured different filament segments in focus. Figure 3A shows the MIF image, the image slice in the image stack with the highest edge noise gradient resulting from the maximum amount of filament area in focus. It can be observed that multiple filaments were partially or fully out-of-focus. Using a computer vision algorithm, AIF images were created from the image stack. This algorithm was able to resolve both small particles and large filaments, all in one single image (Figure 3B). The AIF image successfully reconstructed the full length of filaments; however, it exhibited increased noise relative to the MIF image, with a measured noise level of 2.1 compared to 0.55 for MIF, where larger values indicate greater image noise and less desirable. Consistently, the Laplacian variance was unusually higher for the AIF image (316) than for MIF (9.9), indicating elevated high-frequency content associated with noise.

**Figure 3:**
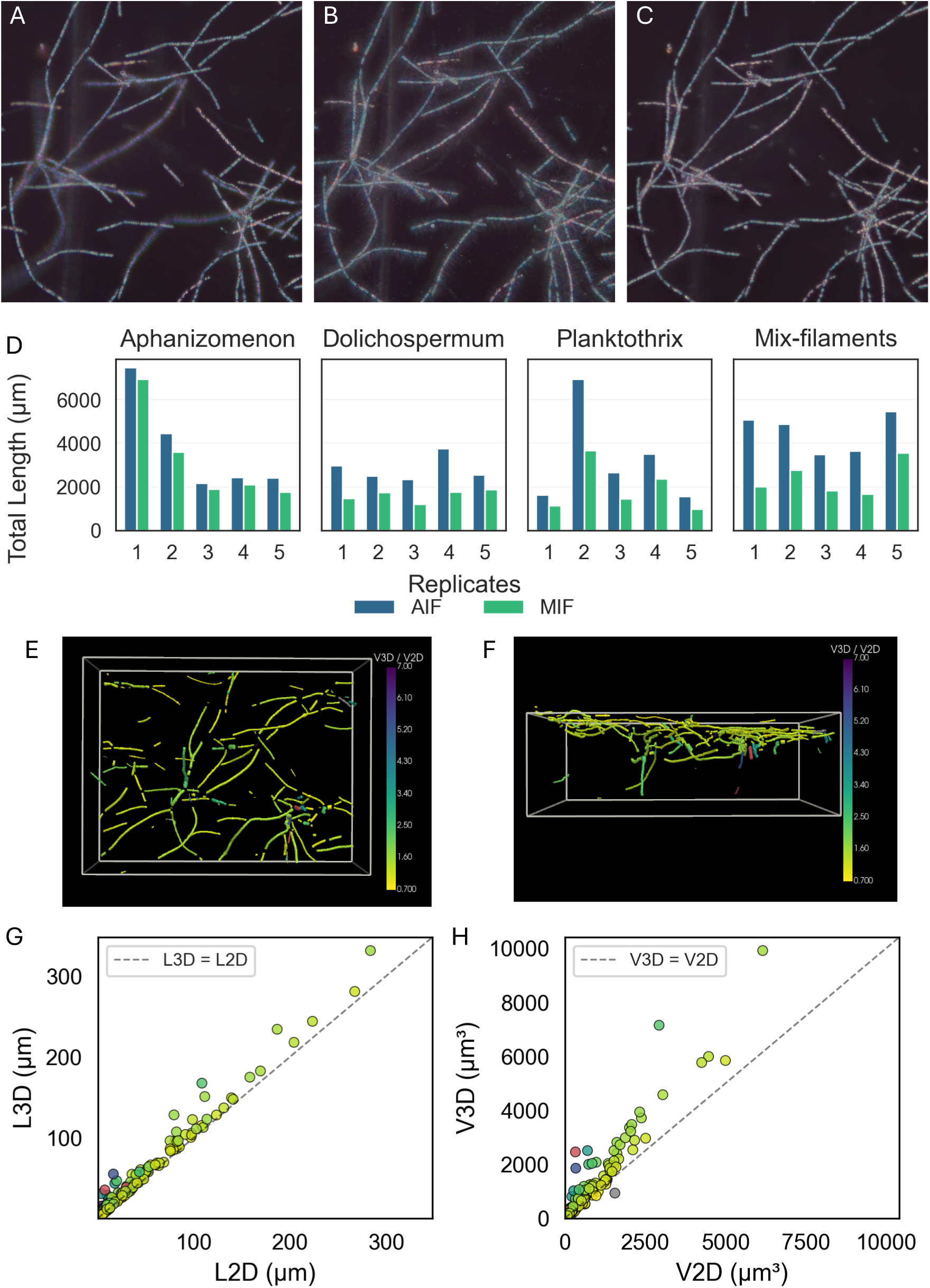
(A) Representative most-in-focus (MIF) frame extracted from a volumetric scan.(B) All-in-focus (AIF) image generated by combining information across multiple focal planes. (C) Denoised AIF image produced using a deep learning–based approach. (D) Total filament length quantified from AIF, MIF, and single-plane imaging (SPI) for three filamentous cyanobacteria: *Aphanizomenon*, *Dolichospermum*, and *Planktothrix*; as well as a mixed sample containing all three species. Each bar cluster represents replicate measurements, with three bars corresponding to AIF, MIF, and SPI. (E-F) Top (E) and side (F) views of reconstructed three-dimensional (3D) filament structures within the imaging volume, colored by ratio of volume measure from 3D reconstruction (V3D) and volume measured from 2D projection (V2D). Filaments with extreme ratios (V3D/V2D<0.7 and V3D/V2D>7) are displayed in gray and red respectively. (G–H) Scatter plots comparing filament length (G) and volume (H) measured from 3D reconstructions versus 2D projections, with points colored according to the same V3D/V2D ratio convention as the 3D plots.

To mitigate this, the AIF image was processed using a deep learning-based denoising model (Figure 3C). The denoised output preserved filament continuity while reducing noise (0.28) and Laplacian variance (5.6) to levels comparable to the MIF image. Although denoising resulted in a moderate reduction in edge sharpness, as reflected by the lower Laplacian variance relative to the MIF image (5.6 vs. 9.9), structural similarity analysis demonstrated substantially improved agreement with the MIF reference image. The denoised image achieved a Structural Similarity Index Measure (SSIM) of 0.947, compared to 0.80 for the original AIF image, indicating enhanced preservation of underlying filament structure and overall image fidelity. Collectively, these results suggest that deep learning-based denoising provides an effective balance between noise suppression and structural preservation in AIF reconstructions.

A custom masking method (see Materials and Methods) was developed to calculate the area of filaments (Figure S6B). The mask was skeletonized and pruned to measure the length of filaments (Figure S6C). The measured lengths should not be interpreted as absolute values, as the skeletonization algorithm does not accurately preserve filament centerlines at locations where foreground masks from multiple filaments intersect. Nevertheless, this approach provides a consistent framework for comparative analysis between AIF and MIF representations acquired from the same field of view. The calculated total area and length of filaments measured from the AIF were 10840 µm^2^ and 5111.7, respectively. An area of 6420.5 µm^2^ of filaments was in focus in the MIF, which is 59% of the total area detected in AIF. Total resolvable length in MIF was 3193 µm (62% of AIF-derived total length). Thus, volumetric scan-derived AIF proved to be a suitable method for analyzing filamentous algae and cyanobacteria.

The resolving capability of AIF images was evaluated for multiple filamentous cyanobacterial species. AIF images were benchmarked against MIF images. Imaging datasets were acquired for *Aphanizomenon sp., Dolichospermum sp*., *Planktothrix agardhii*, and a mixed sample containing all three filamentous taxa. Performance was evaluated by total detected filament length as an indicator of structural resolvability. Figure 3D presents the total detected filament length for five different fields of view for each sample type. AIF consistently yielded a higher total filament length than the MIF. The *Aphanizomenon* filaments often formed entangled networks that were predominantly distributed within a single focal plane. Consequently, a substantial fraction of the filament structures could be resolved near the optimal focal depth, resulting in a significantly greater mean total detected length (3264 µm) in MIF. However, portions of the filaments remained distributed outside this single focal plane and could not be resolved fully. In comparison, AIF integrates information across multiple focal planes, resulting in a higher mean total filament length (3790 µm, MIF/AIF = 0.84).

The difference between AIF and MIF was particularly pronounced for *Dolichospermum* and *Planktothrix*, where mean total filament lengths measured using AIF image were 2828 µm and 3262 µm, respectively, compared to only 1617 µm and 1921 µm measured using MIF image. *Dolichospermum* filaments were short in length, stratified mainly into two layers. MIF only contained particle information for the filaments from the focus layer. AIF resolved particles from both layers, resulting in an approximately twofold (MIF/AIF = 0.58) increase in the mean total detected filament length. *Planktothrix* filaments were long and largely stratified into multiple discrete layers. They did not form entangled three-dimensional structures like *Aphanizomenon*. On the contrary, they were not short like *Dolichospermum* and thus could not be fully resolved from a single plane. Due to the extended length and multi-layer distribution of the filaments, MIF could not completely resolve individual structures. As a result, MIF underestimated the total filament length (MIF/AIF = 0.61). The mixed-filament sample exhibited a similar trend, with AIF resolving almost twice the filament length (MIF/AIF = 0.52). AIF image enables reconstruction of the two-dimensional (2D) projection of filaments distributed across multiple focal planes in one single frame. However, filamentous structures are not strictly confined to a single x-y plane and often exhibit three-dimensional (3D) orientations within the imaging volume. While AIF can provide a good estimation of length and area in the 2D projection, it does not capture the true spatial geometry of filaments, potentially leading to underestimation of filament length and total biovolume. In principle, volumetric scans can provide three-dimensional structural information for filamentous particles, aiding in correct length and biovolume estimation (Figure S7). However, precise axial quantification requires an optical system with a depth of field shallower than the desired z-resolution and effective rejection of out-of-plane light, as achieved in confocal microscopy. ARTiMiS VFI optical configuration provides neither out-of-plane light ray rejection nor sufficiently shallow depth of field. Consequently, direct 3D reconstruction from the volumetric scan data produces artificially enlarged dimensions along the z-axis (Figure S8). To mitigate this issue, 3D construction of filaments was carried out using estimated filament width and z-axis position of each segment of the filaments and considering filaments as a cylindrical shape (Figure 3E and F). The x-y projection of the reconstructed volume (Figure 3E) closely resembled the AIF representation; however, the reconstructed 3D model additionally enabled visualization in the x-z plane (Figure 3F), providing complementary perspectives of the spatial organization and orientation of filaments within the sample matrix. These 3D representations conserve the intrinsic size and orientation of the filaments in 3D space. Filament length and volume were measured from the 3D representations and compared to length and volume measured from AIF (Figure 3G and H). Validation of the 3D-derived filament length and volume measurements was limited to synthetic datasets, as ground-truth measurements for real filamentous samples were unavailable. Consequently, further validation using experimentally controlled filament structures will be necessary in future studies. Nevertheless, 3D-derived length and volume were mostly similar to 2D-derived length and volume for filaments that were oriented flat in the x-y plane. Filaments that were inclined with respect to the z-axis (Figure 3F), 3D-derived length and volume were higher than those of the 2D (color coded by the ratio), providing indirect support for the validity of the reconstruction approach. Total biovolume estimated from 3D was 1.71×10^5^ µm^3,^ which was 47% greater than the biovolume estimated from 2D (1.16×10^5^ µm^3^). Similarly, the total length estimated from 3D was 8109 µm, which was 16.8% greater than the length estimated from 2D (6945 µm). The observed increases in estimated filament length and biovolume demonstrate the potential of VFI-derived 3D reconstruction to overcome limitations associated with 2D projection-based characterization.

### VFI enables mass density estimation across standardized particles and cellular populations

From the previous sections, we have established that ARTiMiS VFI can measure the size and the z-axis position of the particles with high accuracy. This capability unlocks an entirely new class of measurements that have traditionally been difficult to obtain in flow-imaging systems: particle-specific sinking velocity and density. Sinking velocity of three particle types characterized by different diameters and densities: 4.5 µm (1.05 g/cm^3^), 4.8 µm (1.07 g/ cm^3^), and 4.0 µm (1.22 g/cm^3^), was measured using ARTiMiS VFI. Figure 4A presents the distribution of measured sinking velocity (m/s), on the order of 10⁻⁶ m/s. The blue dotted lines represent the theoretical sinking velocity calculated using Stokes’ law. The 4.8 µm particles exhibited strong agreement between experimentally measured (0.875 × 10⁻⁶ m/s) and theoretical sinking velocity (0.876 × 10⁻⁶ m/s). In contrast, the 4.5 µm microspheres exhibited higher median sinking velocity (0.66 × 10⁻⁶ m/s) relative to the theoretical value (0.55 × 10⁻⁶ m/s) with a standard deviation of 0.165 × 10⁻⁶ m/s. This variability is likely associated with the measurement sensitivity of the system. At a z-step size of 10 µm, the average velocity sensitivity of the instrument was estimated to be approximately 0.21 × 10⁻⁶ m/s across the range of possible particle travel distances, placing the observed standard deviation within the expected sensitivity range of the instrument. Although reducing the z-step size could further improve instrument sensitivity, the minimum achievable step size in the current configuration is constrained by the balance between particle settling dynamics, objective lens translation speed, and the time required for image acquisition during volumetric scanning. 4 µm particles exhibited a bimodal sinking velocity distribution, with one peak between 1.25 × 10⁻⁶ and 1.35 × 10⁻⁶ m/s and a second peak between 1.9 × 10⁻⁶ and 1.95 × 10⁻⁶ m/s. The lower-velocity population likely originated from particles that reach the bottom surface before sufficient settling displacement could occur between consecutive scans, thereby reducing the measurable travel distance. To remove measurements associated with the lower-velocity population, a filtering threshold of 1.5 × 10⁻⁶ m/s was applied. The median sinking velocity of the filtered population (1.955 × 10⁻⁶ m/s) closely matched the theoretical prediction (1.91 × 10⁻⁶ m/s).

**Figure 4:**
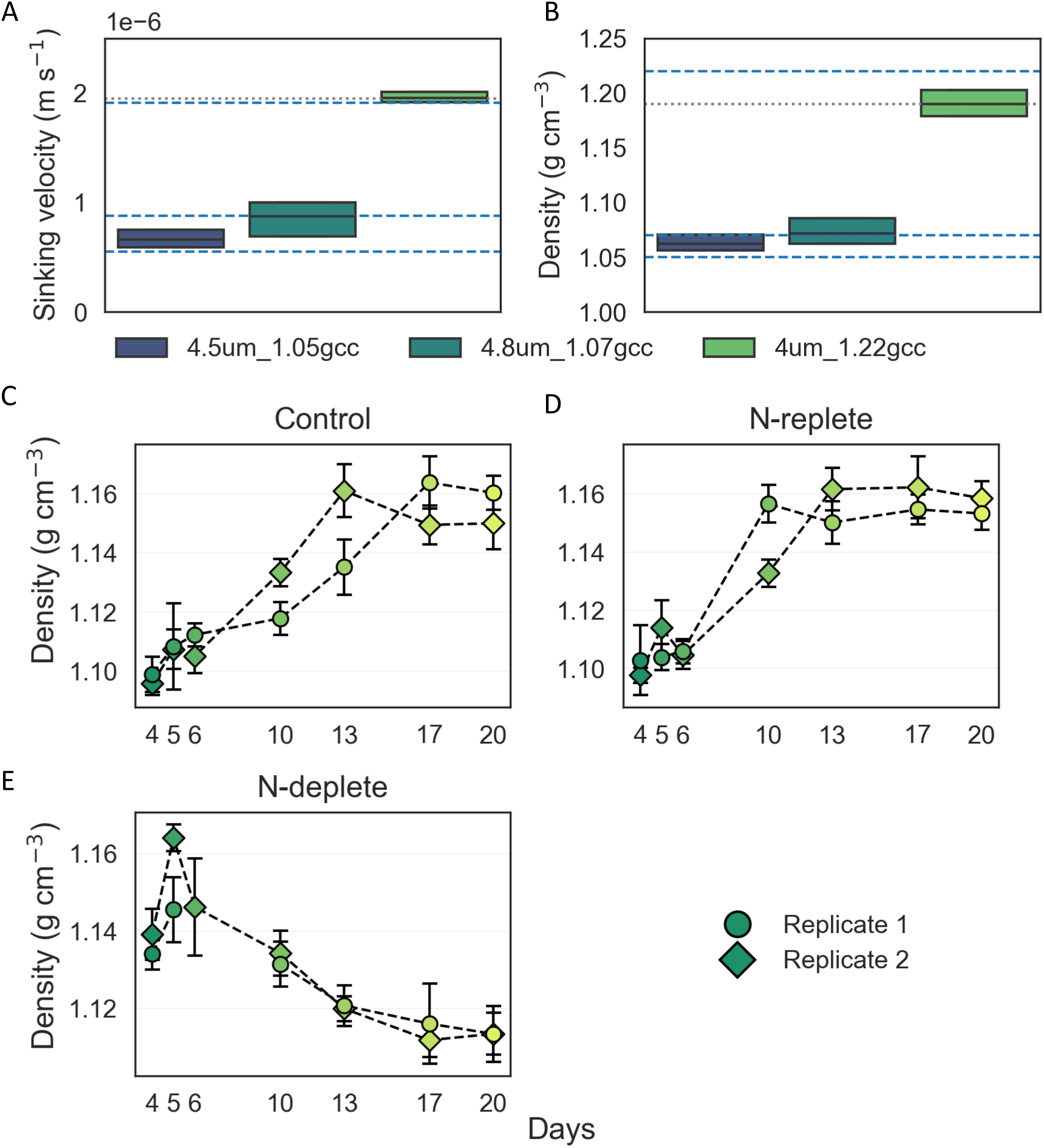
(A–B) Sinking velocity (A) and mass density (B) of microspheres with diameters in the 4–5 µm range and varying densities (1.05, 1.07, and 1.22 g cm⁻³) measured using ARTiMiS. The blue dotted lines indicate theoretical predictions of sinking velocity and density. The gray dotted line in (A) represents the theoretical sinking velocity of 4 µm, 1.22 g cm⁻³ microspheres calculated using an effective diameter of 4.35 µm, while in (B) it represents the corresponding theoretical density based on NIST reference values for the microsphere material. (C–E) Mean mass density of *C. vulgaris* cells in control (C1 and C2), N-replete (NR1 and NR 2) and N-deplete (ND1 and ND2) cultures over the cultivation period. Error bars denote variability across replicate measurements.

As the VFI enables precise measurement of particle size and sinking velocity, we can utilize equation 3 to estimate the mass density of each particle. Figure 4B shows estimated mass density (g/cm^3^) for the three types of microspheres. The measured mass density of 4.8 µm microspheres was fully in agreement with the specification of the microspheres (mean 1.075 g/cm^3^, median 1.071 g/cm^3,^ and a standard deviation of 0.027 g/cm^3^). For 4.5 µm, 1.05 g/cm^3^ microspheres, ARTiMiS derived mass density (mean 1.064 g/cm^3^, median 1.062 g/cm^3^) was higher than the manufacturer’s specified value (1.05 g/cm^3^). This overestimation is consistent with the measured sinking velocity, which was also higher than the theoretical value estimated by Stokes’ law. As settling velocity scales directly with the density difference between the particle and the fluid, a higher measured sinking velocity corresponds to a higher inferred particle density. A greater discrepancy was observed between the measured and the specified mass density for 4 µm microspheres. The manufacturer specified density of the polymethyl methacrylate (PMMA) microspheres was 1.22 g/cm^3^, whereas ARTiMiS VFI reported value was 1.190 g/cm^3^, which is the same as the mass density of PMMA according to the National Institute of Standards and Technology (NIST). Upon further inspection, we determined that the particle size distribution for these microspheres significantly differed from the specified value. The size distribution reported by ARTiMiS VFI indicates a mean particle diameter of approximately 4.35 µm and an area of approximately ∼14.7 µm^2^ (Figure S9). This value falls outside the upper bound of the manufacturer’s specification (4.00 ± 0.15 µm; coefficient of variation < 3%). Using the ARTiMiS VFI-derived mean diameter of 4.35 µm and a specific gravity of 1.19, the theoretical sinking velocity calculated from Stokes’ law (equation 2) was 1.95 × 10⁻⁶ m/s, which closely matched the experimental value (1.955 × 10⁻⁶ m/s). Furthermore, when mass density was recalculated using the ARTiMiS-derived particle diameter and measured sinking velocity, the mode of the resulting density distribution was 1.190 g/cm^3,^ consistent with the NIST-reported value for the material. This agreement suggests that the apparent discrepancy with the catalog-specified density may stem from the nominal diameter assumption of 4 µm. To demonstrate that the mass density measurement is size agnostic, 6 µm microspheres with a density of 1.05 g/cm^3^ were also analyzed with ARTiMiS VFI. Measured sinking velocity and mass density were slightly higher than the theoretical value, similar to 4.5 µm beads (Figure S10). These results demonstrate that ARTiMiS VFI can measure particle sinking velocity and estimate the mass density of spherical particles.

This framework was then extended to measuring the effect of nitrogen starvation on *C. vulgaris* cultured under controlled laboratory conditions. Cell size and mass density changes in *C. vulgaris* culture due to nutrient variation in culture media. Nitrogen deprivation induces an increase in cell size and lipid content, accompanied by a decrease in mass density (Baroni et al., 2019; Liu et al., 2022; Zhang et al., 2013). *C. vulgaris* cells were grown in two flasks (C1 and C2) in Bold 1NV media as controls. Two additional cultures (NR1 and NR2) were established by collecting cells from the two flasks, mixing and distributing them in Bold 1NV media containing flasks. These four flasks should show similar trends in cell phenotype expression. Figure 4C-D shows the changes in mean cell mass density during the 20-day culture period. All four cultures had low cell mass density during the initial days (day 4-6) of the culture and high density at the later days (day 13-20). NR flasks reached the stable density earlier than the control flasks (day 10-13 vs day 13-17). For all four flasks, mass density increased from ∼ 1.1 g/cm^3^ to ∼ 1.16 g/cm^3^. This shows the batch effect on *C. vulgaris* cells. As cell concentration increases (Figure S11) and nutrient depletes, *C. vulgaris* cells go from low mass density to high mass density. ARTiMiS VFI was sensitive enough to detect small changes in mass density in the cell population.

To see the effect of nitrogen depletion in the culture, two additional flasks (ND1 and ND2) with modified Bold 1NV media (no nitrogen) were inoculated using the cells from the cell mix from C1 and C2. In nitrogen deprivation, *C. vulgaris* cells become growth-limiting, increase cell size, and start accumulating lipid and other macromolecules in nitrogen-limited conditions, resulting in changes in mass density and buoyancy. Previous studies have reported that nitrogen deprivation induces cell mass density reduction and lipid accumulation (Chu et al., 2020; Jiang et al., 2019). Cells in ND1 and ND2 reached ∼1.145 g/cm^3^ density on day 6, and then it continuously decreased till the end of the culture period (Figure 4E). This trend was opposite to that of the control and NR flasks. Baroni et al. (2019) reported similar trends in *C. vulgaris* batch cultures, where the cell density initially increased before progressively decreasing over the remainder of the cultivation period. Although the experimental conditions differed between the two studies, including the *C. vulgaris* strain and the density measurement approach, the trends in cell size and density observed in the present study are consistent with the findings of Baroni et al. (2019), further supporting the physiological interpretation of the measurements. These findings indicate that ARTIMIS VFI can not only precisely measure culture concentrations and individual cell sizes, but also measure individual cell mass density, all using one single instrument. Importantly, the integration of these different dimensions of information can be directly utilized to manage and adapt maintenance, scale-up, and harvest parameters to optimize algae-based product yield.

## Conclusion

This study demonstrated that VFI can overcome key limitations of FIM by enabling improved particle resolvability, enhanced image quality, and advanced particle characterization. VFI scans through the depth of the channel and extracts particle images from the optical section corresponding to the particle’s axial position, thereby enabling near-optimal focal representation of particles distributed throughout the imaging volume. This increases particle detection accuracy, reduces out-of-focus particles, provides sharper particle image crops, and thus enables precise particle characterization. Image stacks acquired through VFI enable AIF image generation, which not only enables simultaneous visualization of particles distributed across multiple x-y planes, but also enables imaging of long filamentous particles extending across multiple planes in the z-axis that are difficult to fully resolve using conventional single plane imaging. VFI can further extend the analytical capability of FIM technology. In addition to improving image quality and analytical precision, VFI makes it possible to measure dynamic properties, including sinking velocity and mass density. Beyond accurately estimating the mass density of standard microspheres, VFI also captured temporal trends in the mass density of *C. vulgaris* cells under nitrogen-replete and nitrogen-deplete conditions. These capabilities highlight the potential of VFI as a powerful tool for monitoring physiological and biophysical changes in microalgal cultures. Collectively, these results indicate VFI as a promising framework for the next generation of FIM, offering improved precision in population characterization.

## Acknowledgments

This research was supported by the U.S. Department of Energy, Office of Energy Efficiency and Renewable Energy (EERE), under Award Number DE-EE0011058, the Georgia Research Alliance (GRA), and the Georgia Artificial Intelligence in Manufacturing (GA-AIM) initiative.

