## Supplemental Information for "Volumetric Flow Imaging Microscopy to Enhance Particle Characterization"

### Supplementary Information

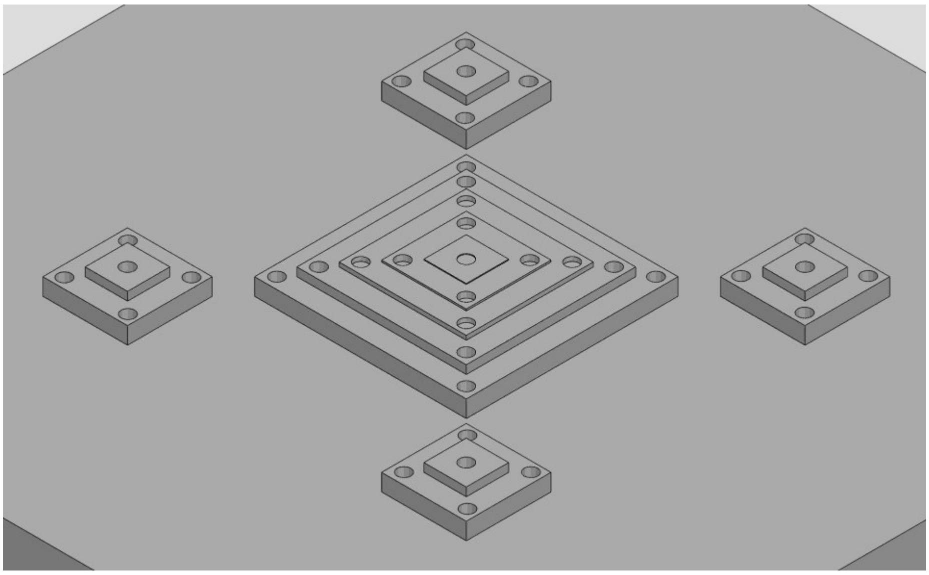

Figure S1: Step pyramid used to calibrate the voice coil motor. The structure has one central pyramid and four peripheral pyramids. Central pyramid contains five steps (20 µm, 10 µm, 5 µm, 2 µm, and 1 µm) and used for step size calibration. Peripheral pyramids have two steps (20 µm and 10 µm) and used for leveling optical assembly.

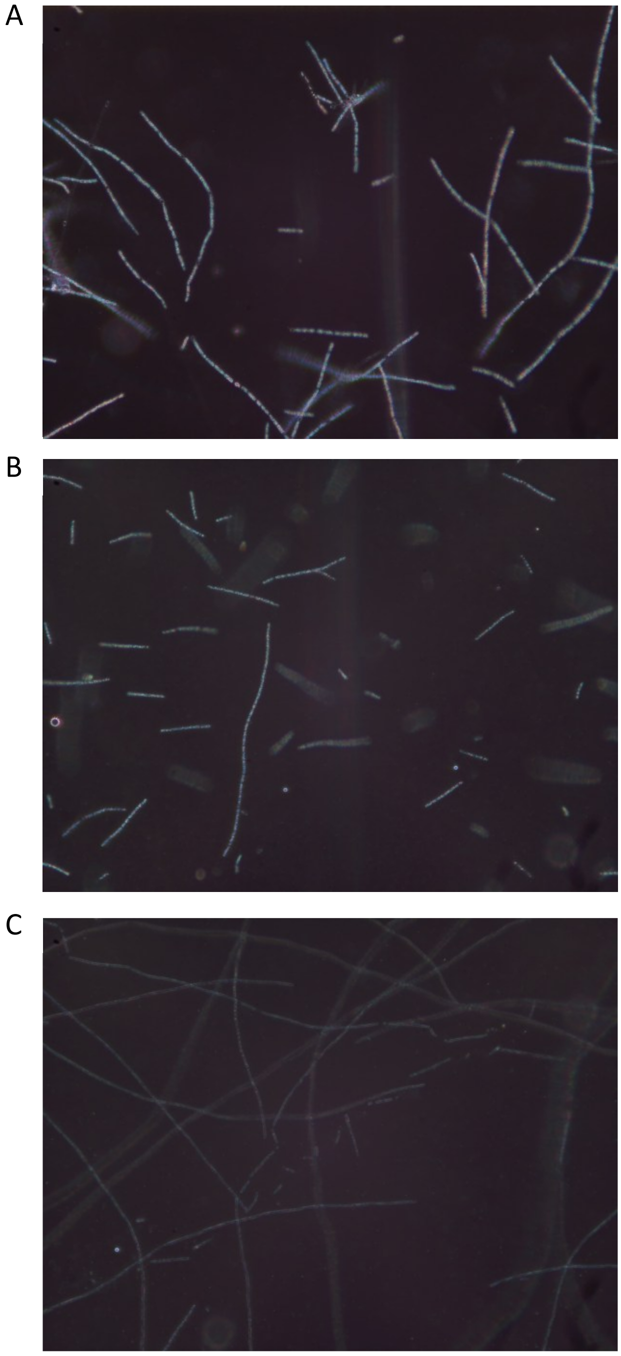


Figure S2: Darkfield images of (A) *Aphanizomenon sp.*, (B) *Dolichospermum flos-aquae,* and (C) *Planktothrix agardhii*

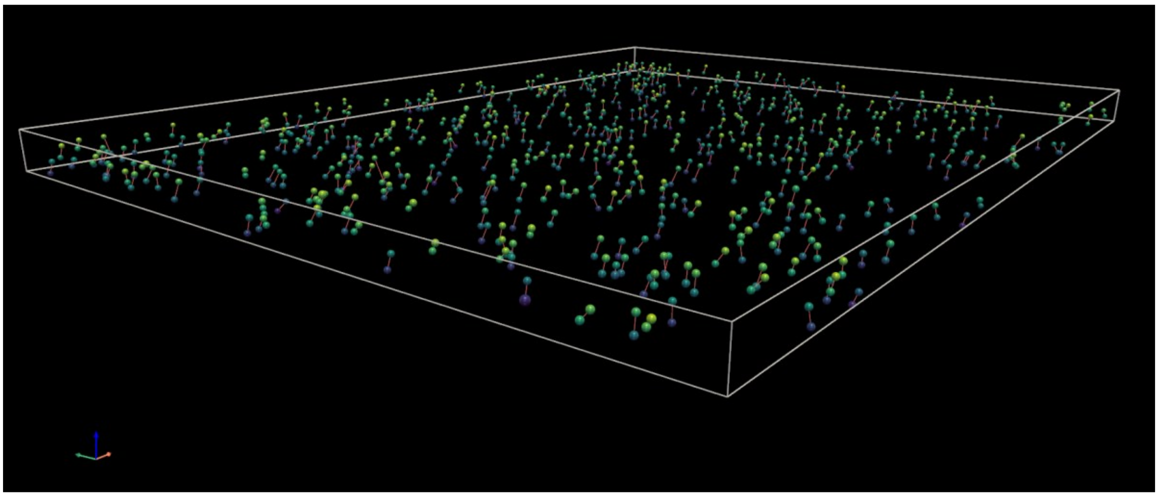

Figure S3: Demonstration of bipartite matching of particles from two consecutive volumetric scan. Matched particles are connected with a red line.

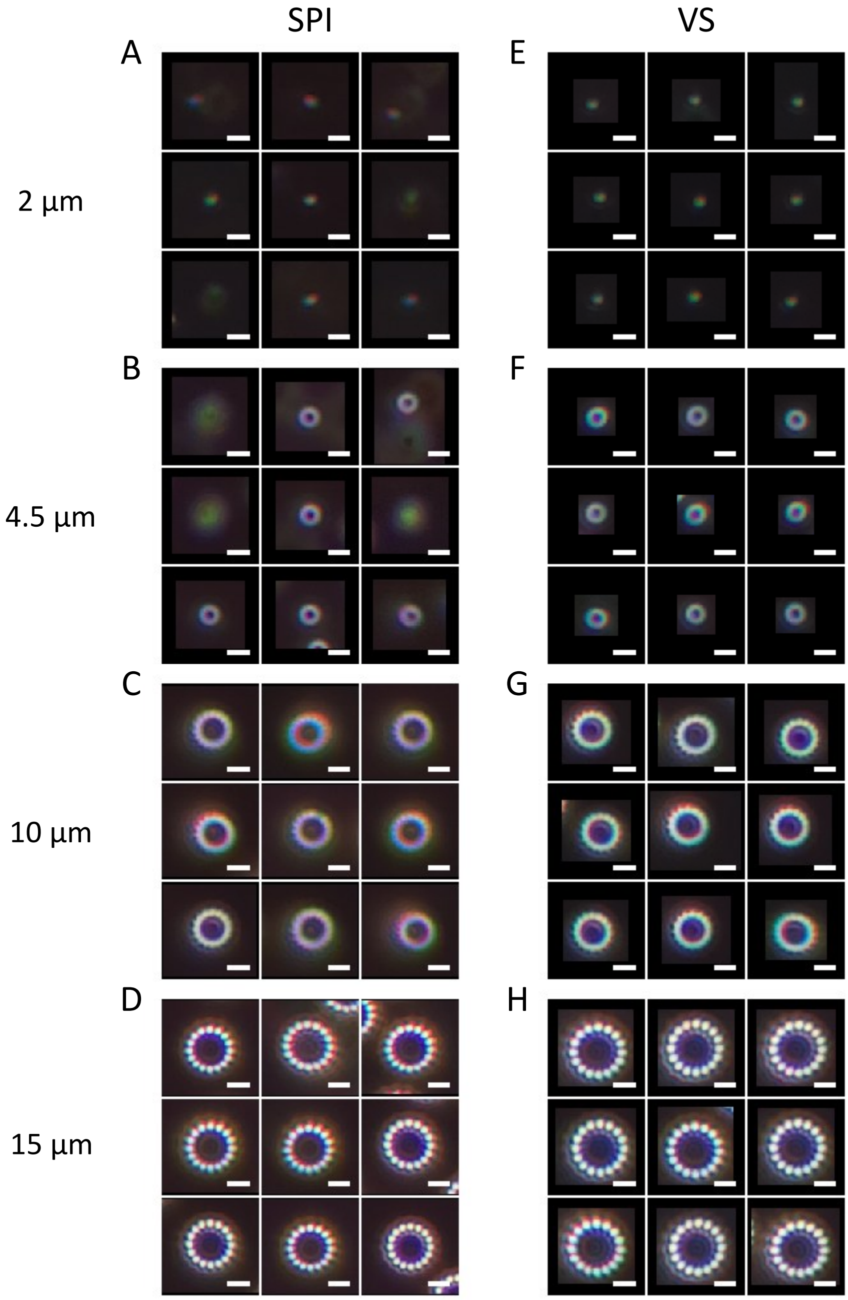


Figure S4: Representative ROIs for 2, 4.5, 10 and 15 µm microspheres captured using SPI and VS method. A portion of the ROIs derived from SPI were blurry and out of focus while all ROIs from VFI were sharp and in focus.

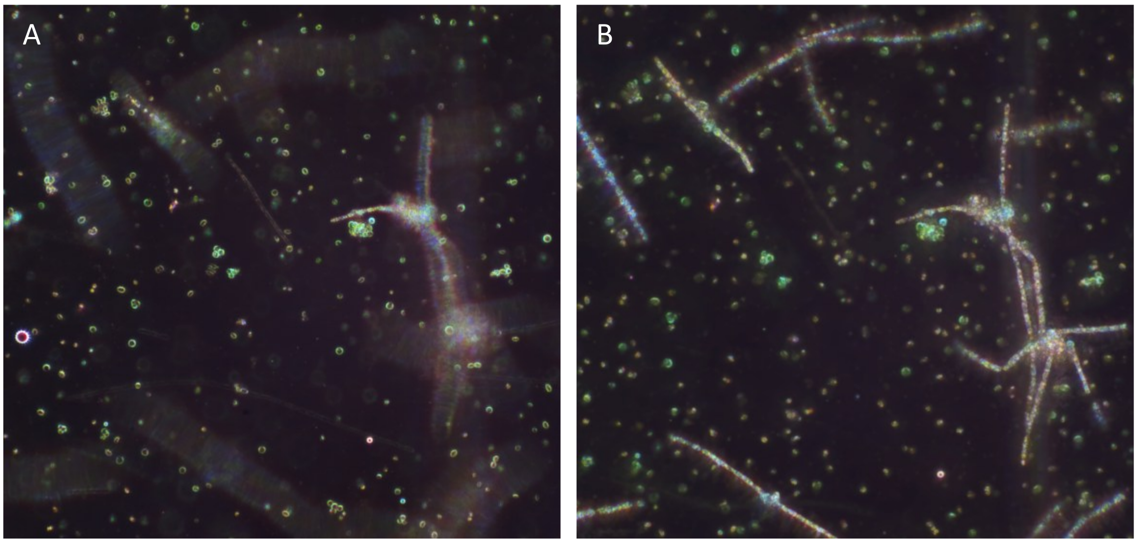


Figure S5: (A) Single plane image of a sample matrix containing spherical cells, colonies, and filaments. (B) AIF image generated from the volumetric image stack acquired using VFI. AIF image can provide a comprehensive representation of the sample matrix.

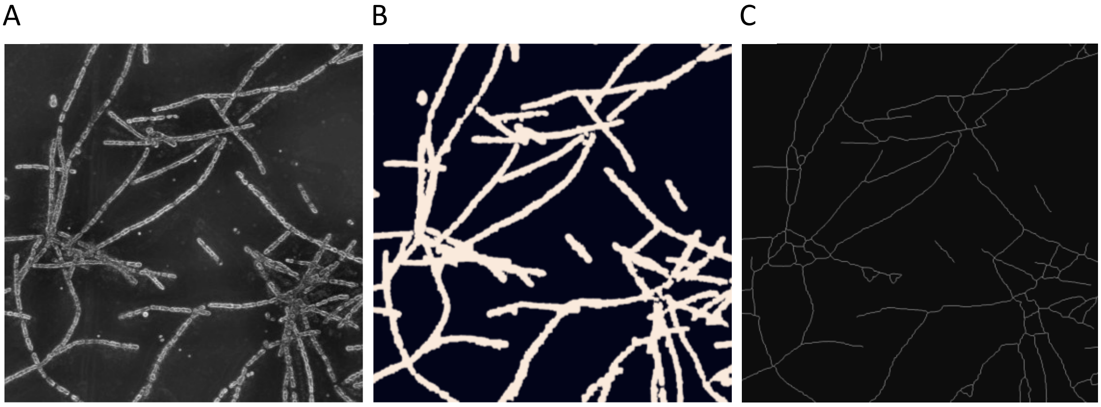


Figure S6: (A) Gradient map of all in focus image generated from volumetric scan of *Aphanizomenon* sample, (B) corresponding binary mask separating the filaments (foreground) from the background used for filament area calculation, (C) Pruned skeleton of binary mask depicting the centerline of the filaments used to calculate length of the detected filaments.

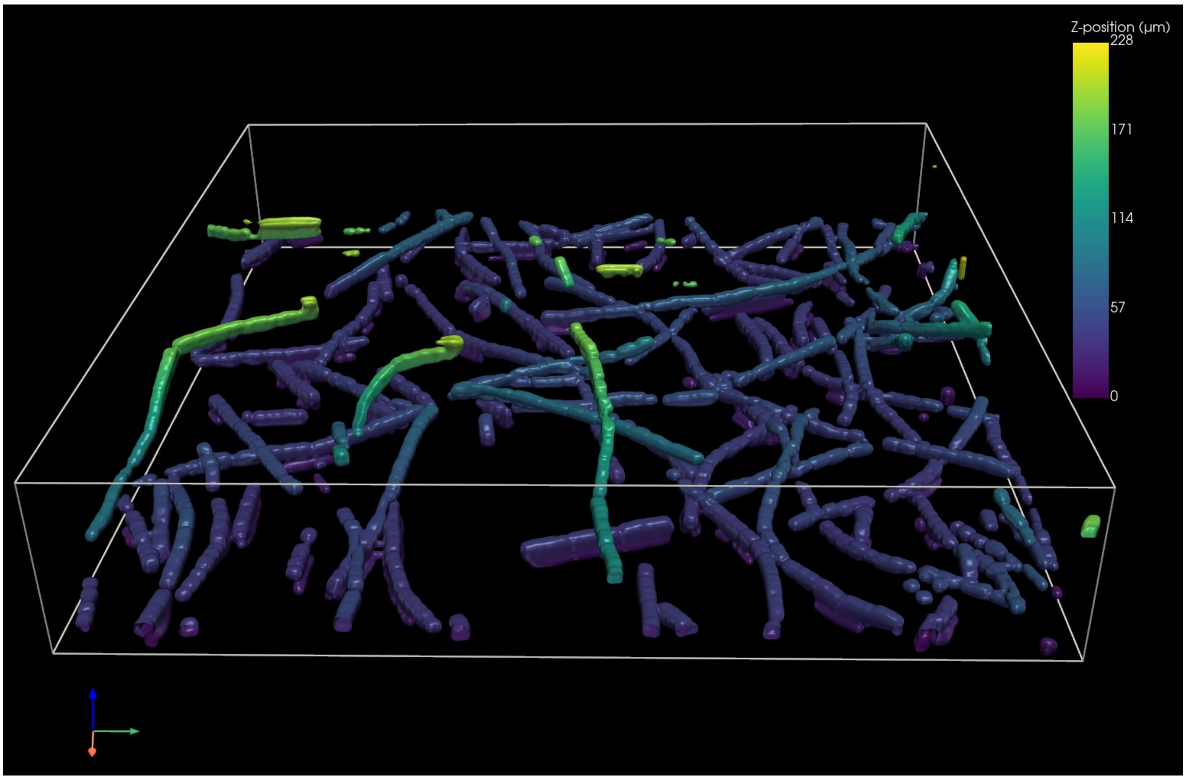


Figure S7: Isometric view of reconstructed three-dimensional (3D) filament structures directly from the volumetric scan captured with Nikon W1 spinning disk confocal microscope with 60x oil objective and a step size of 1 μm.

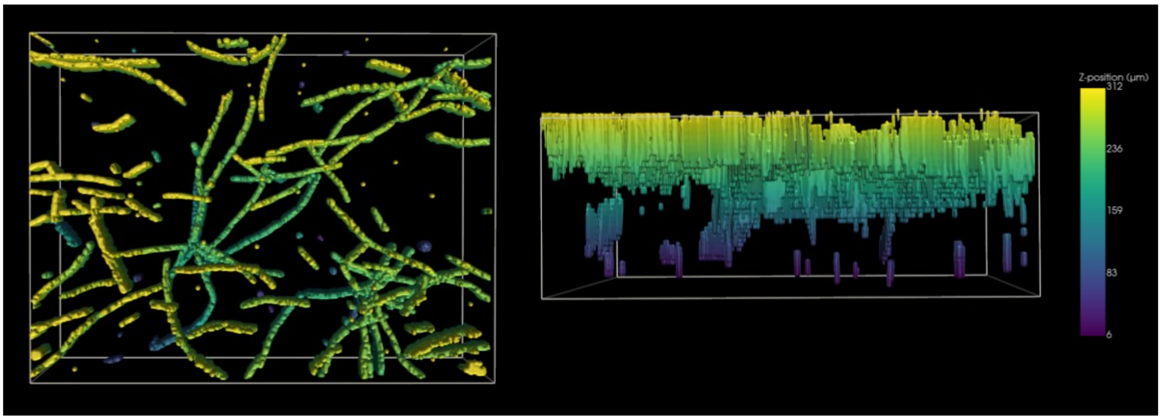


Figure S8: Top and side views of reconstructed three-dimensional (3D) filament structures directly from the volumetric scan with a step size of 10 μm within the imaging volume, colored by axial (z-axis) position.

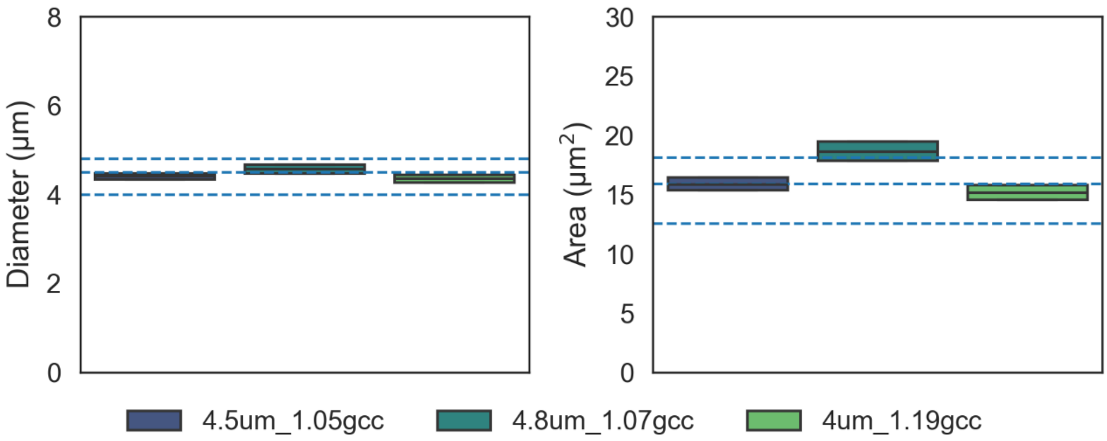


Figure S9: Box plot of microsphere diameters (left) and area (right) measured using ARTiMiS. Blue dotted lines represent manufacturer recommended diameter and theoretical area computed from the nominal diameter.

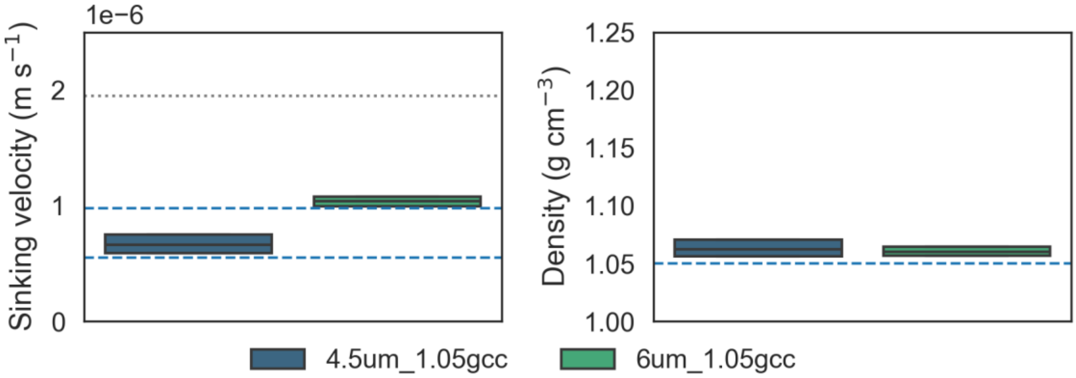


Figure S10: Box plot of sinking velocity (left) and mass density (right) of 4.5 and 6 μm microspheres with a nominal mass density of 1.05 gm cm$^{-3}$ measured using ARTiMiS VS. Blue dotted lines represent theoretical sinking velocity and nominal mass density.

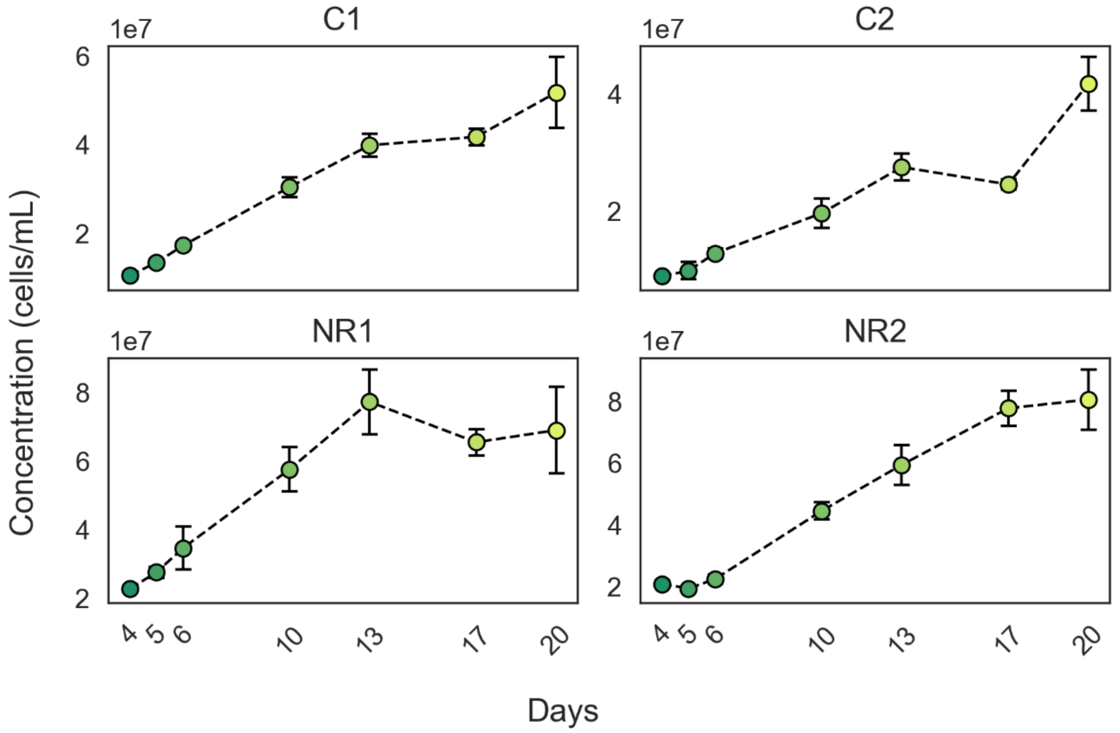


Figure S11: Cell concentration in C1, C2, NR1, and NR2 cultures over the cultivation period.
